# Enhanced Peripheral Nerve Regeneration by Mechano-electrical Stimulation

**DOI:** 10.1101/2023.04.20.537728

**Authors:** Youyi Tai, Thamidul Islam Tonmoy, Shwe Win, Natasha T. Brinkley, B. Hyle Park, Jin Nam

## Abstract

To address limitations in current approaches for treating large peripheral nerve defects, this study evaluated the efficacy of functional material-mediated physical stimuli on peripheral nerve regeneration. Electrospun piezoelectric poly(vinylidene fluoride-trifluoroethylene) nanofibers were utilized to deliver mechanical actuation-activated electrical stimulation to nerve cells/tissues in a non-invasive manner. Using morphologically and piezoelectrically optimized nanofibers for neurite extension and Schwann cell maturation based on in vitro experiments, piezoelectric nerve conduits were implanted in a rat sciatic nerve transection model to bridge a critical-sized sciatic nerve defect (15 mm). A therapeutic shockwave system was utilized to activate the piezoelectric effect of the implanted nerve conduit on demand. The piezoelectric nerve conduit-mediated mechano-electrical stimulation induced enhanced peripheral nerve regeneration, resulting in full axon reconnection with myelin regeneration from the proximal to the distal ends over the critical-sized nerve gap. Furthermore, superior functional recovery was observed by walking track analysis and polarization-sensitive optical coherence tomography, demonstrating the excellent efficacy of the mechano-electrical stimulation strategy for treating peripheral nerve injuries.

## 1. Introduction

Peripheral nerve injury is often associated with severe morbidities such as muscle weakness, loss of sensation, decreased reflexes, and the development of neurological ulcers^1^. Regardless of the origin of the damage, caused by injury or disease, typically the neurotic nerve from significant injuries has to be surgically removed to prevent further damage. Full functional recovery following surgical intervention, however, is difficult to achieve^2, 3^. Potential treatment for nerve injuries with a critical-sized defect is autologous transplantation^4^. Despite advances in surgical technology, complete functional recovery following peripheral nerve damage remains rare, especially for injuries beyond the fourth degree, due to the relatively poor inherent capability of nerves for regeneration^4–6^. Moreover, inevitable denervation from the patient with the autologous surgical procedure results in a loss of function at the donor site^4^. Therefore, severe nerve injury often causes permanent disabilities, devastatingly impacting the quality of life of the patient.

New advanced technology with the use of biocompatible nerve conduits has been reported as an alternative approach for peripheral nerve injury treatments^7, 8^. Biophysical structures as well as various biochemical cues have been incorporated into nerve conduits to facilitate regeneration^9–11^. For example, microgroove patterns of the nerve guidance conduits significantly direct aligned axonal growth, which in turn improves nerve regeneration efficiency^12^. In addition, adding nerve growth factors, especially with a concentration gradient, along the length of the conduits has been proven to further enhance the axonal alignment and extension rate^9^. Despite these improvements, functional recovery is still limited with a critical-sized defect of peripheral nerve injury due to uncontrolled axonal sprouting and the short-lasting effects of the growth factor release at the injury site^13, 14^. Therefore, in addition to the bio-conducive conduit structure and biochemical cues, additional augmenting factors are necessary to promote functional peripheral nerve recovery.

Recently, several studies have recognized the potential of physical stimuli for enhancing regenerative cellular activities, including the application of magnetic fields to augment bone regeneration, the utilization of fluid shear force to promote the vascular formation, and the dynamic control of the substrate stiffness to regulate early-stage pluripotent stem cell differentiation^15–19^. In regard to nerve regeneration, electrical stimulation has been shown to enhance neurite formation and outgrowth both in vitro and in vivo^20, 21^. One of the major challenges in applying electrical stimulation in clinical settings is the need for external electrical devices, which hinders its long-term translational applications^22^.

Piezoelectric materials generate electrical potentials when mechanically actuated^23, 24^. We previously demonstrated that the combination of mechanical actuation, required to induce the piezoelectric effect, and resultant electrical potential from the activation of piezoelectric material synergistically modulate the functionality of neurons and glial cells derived from neural stem cells^25^. In this study, electrospun biocompatible piezoelectric nanofibers were utilized to electrically stimulate peripheral nerve cells, and their effects are examined both in vitro and in vivo. We optimized the electrospun fiber diameter, and the magnitude of electrical stimulation activated by the appropriate mechanical actuation to maximize the regenerative and functional responses from neuronal cells and Schwann cells. Based on these in vitro results, we fabricated piezoelectric nerve guidance conduits to bridge sciatic nerves with a critical-sized defect (15 mm) in a rat model. A shockwave system was used to apply mechanical actuation to the implanted conduits to activate the piezoelectric effect for electrical stimulation. We demonstrate that the mechano-electrical stimulation (MES), derived from the shockwave actuation of the implanted piezoelectric conduits, elicits enhanced peripheral nerve regeneration based on functional motion recovery from the walking track analysis as well as superior axonal re-connection and myelination determined from optical coherence tomography (OCT) and immunohistochemistry. Therefore, this piezoelectric material-based MES strategy provides a means to develop an effective clinical treatment of critical-sized peripheral nerve injuries through non-invasive, long-term physical stimulations.

## 2. Results

### 2.1. Morphologic and piezoelectric optimization of electrospun P(VDF-TrFE) nanofibers

Based on our previous study^25^, electrospun aligned P(VDF-TrFE) scaffolds of various fiber sizes (average diameter of 200, 500, or 800 nm) that present different morphological and piezoelectric characteristics were synthesized (**Figure S1a**). We have shown that electrospun fiber diameter significantly affects neural stem cell behaviors as a larger fiber diameter promotes neurite alignment^25^. In contrast, electrospun P(VDF-TrFE) scaffolds composed of smaller fibers produce greater electric outputs under mechanical perturbation due to their higher piezoelectric constants^25, 26^. To determine the optimal fiber diameter of the electrospun P(VDF-TrFE) nanofibers for peripheral nerve regeneration by balancing the fiber diameter-dependent morphological and piezoelectric characteristics, an in vitro study was conducted to examine the effects of fiber diameter on neuronal cell behaviors. As shown in **Figure 1a, b**, PC12 cells, a neuronal progenitor cell line, exhibited considerably greater alignment on larger fibers (500 nm and 800 nm), as compared to the cells cultured on electrospun P(VDF-TrFE) scaffolds having fibers with an average diameter of 200 nm. In addition, the larger fiber diameters enhanced neurite formation and its extension; there was a significant increase in the number of neurite-bearing cells on 500 nm and 800 nm fiber diameter scaffolds (**Figure 1c**). Furthermore, the average neurite length of the PC12 cells cultured on electrospun P(VDF-TrFE) scaffolds composed of 500 nm- and 800 nm-sized nanofibers was 59 µm and 77 µm, respectively, significantly longer than that of 200 nm at 30 µm (**Figure 1d**). Larger P(VDF-TrFE) nanofibers also induced a greater population of cells that possess high neurite/nucleus ratios (**Figure 1e**). In contrast to these cellular behaviors, larger fiber diameters exhibit smaller piezoelectric constant (d_33_) values (**Figure S1b**). This fiber diameter-dependent piezoelectricity resulted in a greater electric potential generation from electrospun P(VDF-TrFE) scaffolds having smaller fiber diameters under the same mechanical actuation (**Figure S1c**). To ensure the activation of the piezoelectric effect under physiologically safe strain ranges as well as to maximize the cellular alignment and neurite extension, electrospun P(VDF-TrFE) scaffolds having an average fiber diameter of 500 nm were used for the rest of this study.

**Figure 1.**
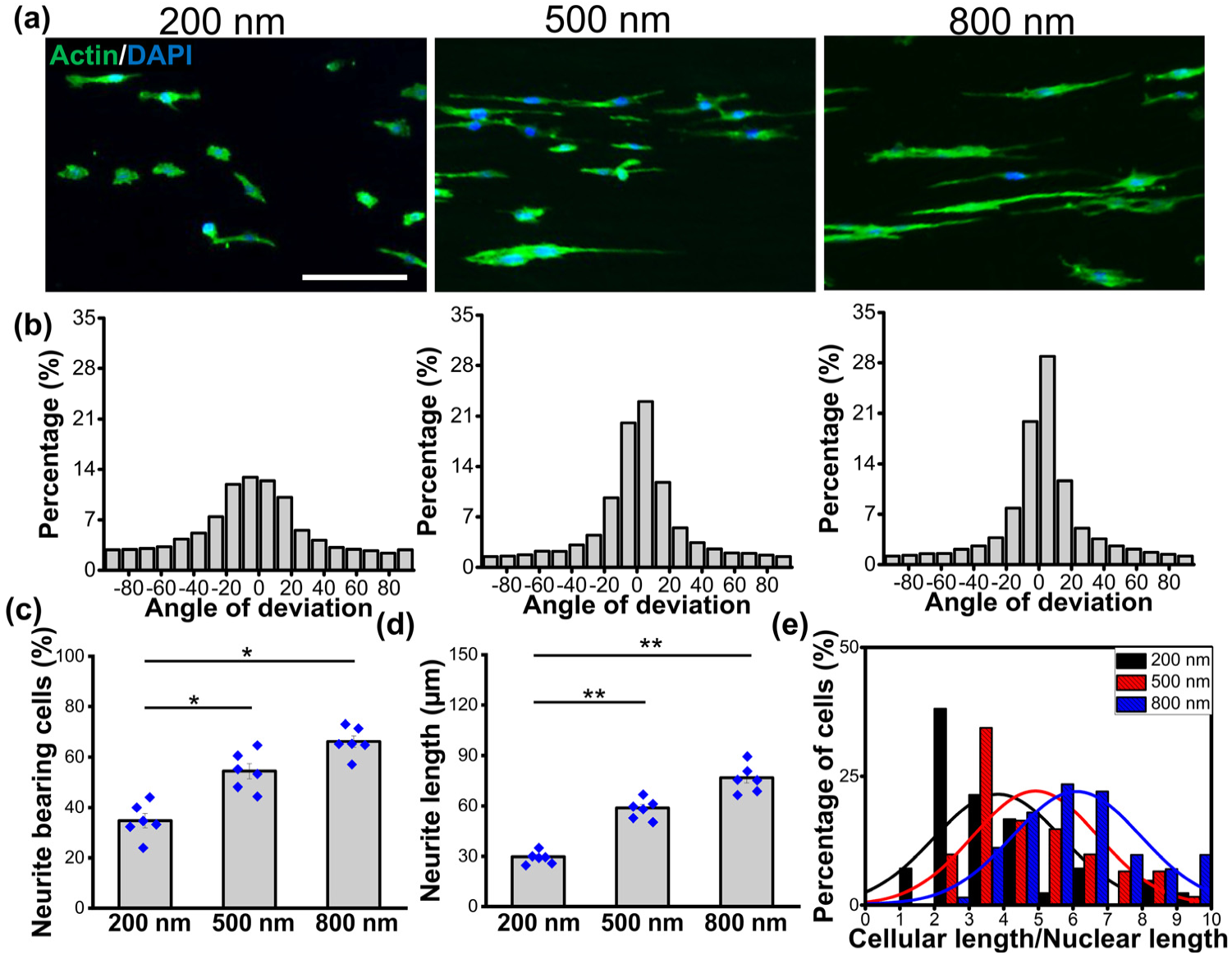
Effects of different fiber diameter on neurite formation and extension. (a) Immunofluorescence images depicting cellular morphology of PC12 cells, which were cultured on electrospun aligned P(VDF-TrFE) nanofibers with average diameters of 200 nm, 500 nm, and 800 nm (scale bar: 100 µm). (b) Directionality histograms of the cells on the scaffolds having various average fiber diameters of (left) 200, (middle) 500, and (right) 800 nm. (c) The percentage of cell population bearing neurites, (d) average neurite length, and (e) the percentage of neurite-bearing cells possessing a certain range of cellular length-to-nuclei length ratio, quantified from immunofluorescence images in (a) (* and ** denote statistical significance of p < 0.05 and p < 0.01, respectively. Six fluorescence images from independent samples were used for each condition).

### 2.2. Optimization of electrical stimulation regimen for neurite formation and extension

To determine the optimal electrical stimulation regimen to promote neurogenesis and neurite extension in PC12 cells, the effects of electric fields with various magnitudes and durations were examined using a custom electrical stimulation device as shown in **Figure 2a**. In addition to the morphological confinement of aligned fibers inducing cellular alignment, electrical stimulation further enhanced neurite formation and neurite outgrowth, which were maximized under the applied voltage magnitude of 200 mV_p-p_ (**Figure 2b**). A notable decrease in cell population under 400 mV_p-p_ stimulation was observed, indicating that a high magnitude of the applied electric field significantly impacts cell viability. A significant increase in neurite-bearing cell population and average neurite length was observed under 200 mV_p-p_ of electrical stimulation, as compared to the control and 100 mV_p-p_ groups (**Figure 2c, d**). Multi-day stimulation (electrical stimulation for 2 hours for 3 consecutive days; ES x 3) did not affect the number of neurite-bearing cells. A substantial increase in the average neurite length was, however, observed under multi-day stimulation of 200 mV_p-p_, as compared to the 1-day stimulation (electrical stimulation for 2 hours, followed by 2 days of rest; ES x 1) (**Figure 2b-d**). It should be noted that the multi-day stimulation induced the greatest number of cells that possess neurites in the range of 6-10 times longer than the nuclear size (**Figure 2e**). Based on these results, a periodic electrical stimulation targeting a magnitude of 200 mV_p-p_ was used for subsequent in vitro and in vivo experiments.

**Figure 2.**
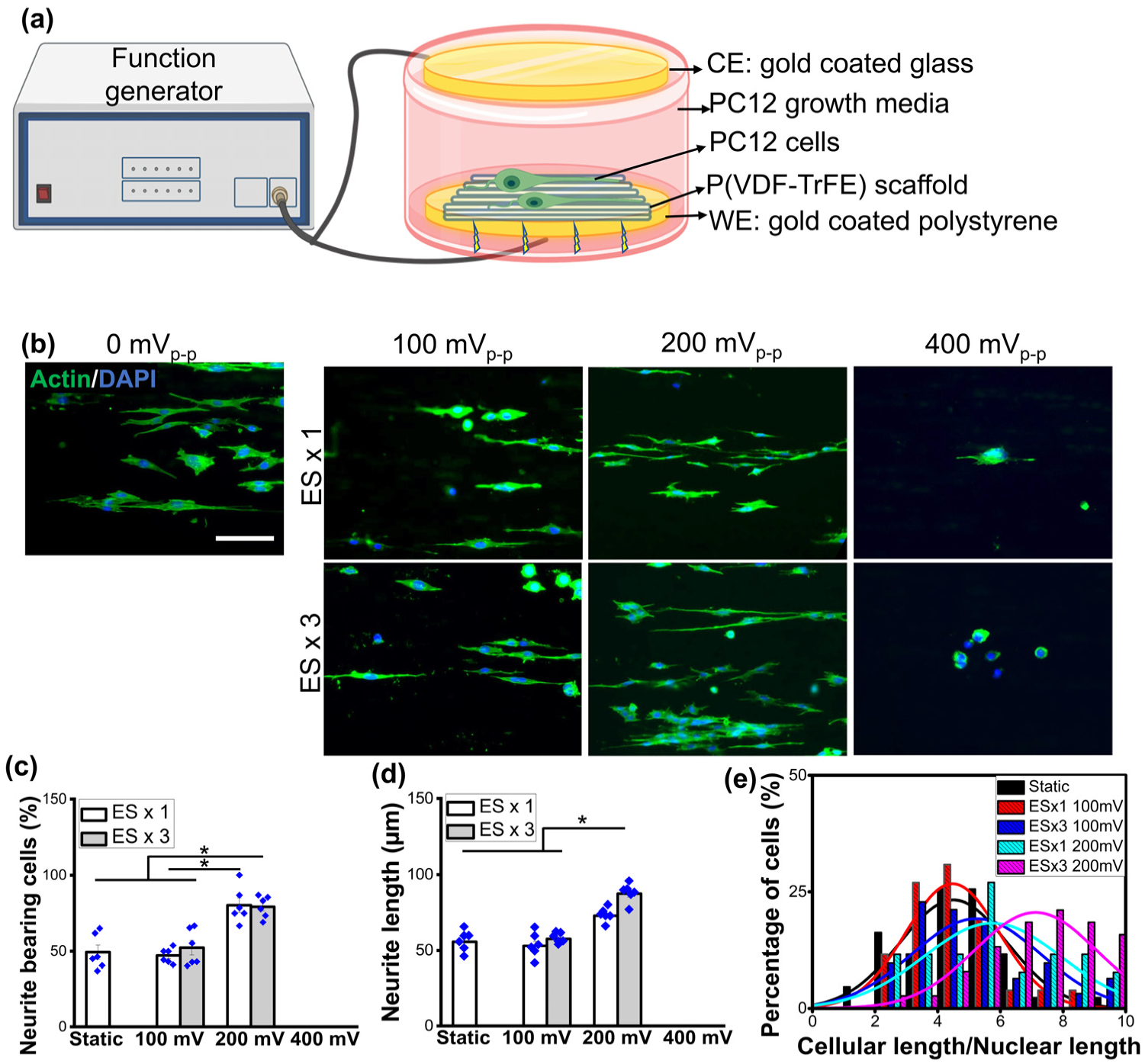
Effects of electrical stimulation (ES) on neurite formation and extension. (a) Schematic of ES setup. (b) Representative immunofluorescence images showing cellular morphology of PC12 cells cultured on P(VDF-TrFE) scaffolds, subjected to a 1-day or 3-day ES with a magnitude of 100, 200, or 400 mV_p-p_ (scale bar: 40 µm). (c) The percentage of cell population bearing neurites, (d) average neurite length, and (e) the percentage of neurite-bearing cells possessing a certain range of cellular length-to-nuclei length ratio, quantified from immunofluorescence images in (a) (* denotes statistical significance of p < 0.05. Six fluorescence images from independent samples were used for each condition).

### 2.3. The effects of mechano-electrical stimulation (MES) on neural cells in vitro

To examine the effects of MES, derived from the mechanical actuation of piezoelectric P(VDF-TrFE) scaffolds^25^, on neuronal cells, PC12 cells were subjected to MES by culturing them on P(VDF-TrFE) scaffolds which were hydro-acoustically actuated by using a customized cell culture system (**Figure 3a**). The thickness of electrospun P(VDF-TrFE) scaffolds having 500 nm-sized fibers was optimized to produce 200 mV_p-p_ under hydro-acoustic actuation to promote pro-neuronal cellular behaviors. A 1-week culture under MES (2 hours per day) significantly enhanced neurite formation and extension as compared to the static culture (**Figure 3b, c**), consistent with the results when electrical stimulation was directly applied, as shown in **Figure 2**. Moreover, the gene expression levels of neuronal markers, *Tubb3* and *Map2*, were significantly upregulated by MES, as compared to the statically cultured cells on the scaffolds (Static group) as well as the control group where cells were cultured on a tissue culture plate (Control group) (**Figure 3d**). A mature neuronal marker, *Chat*, also showed a considerable upregulation under MES, suggesting physical stimulation-mediated neuronal maturation.

**Figure 3.**
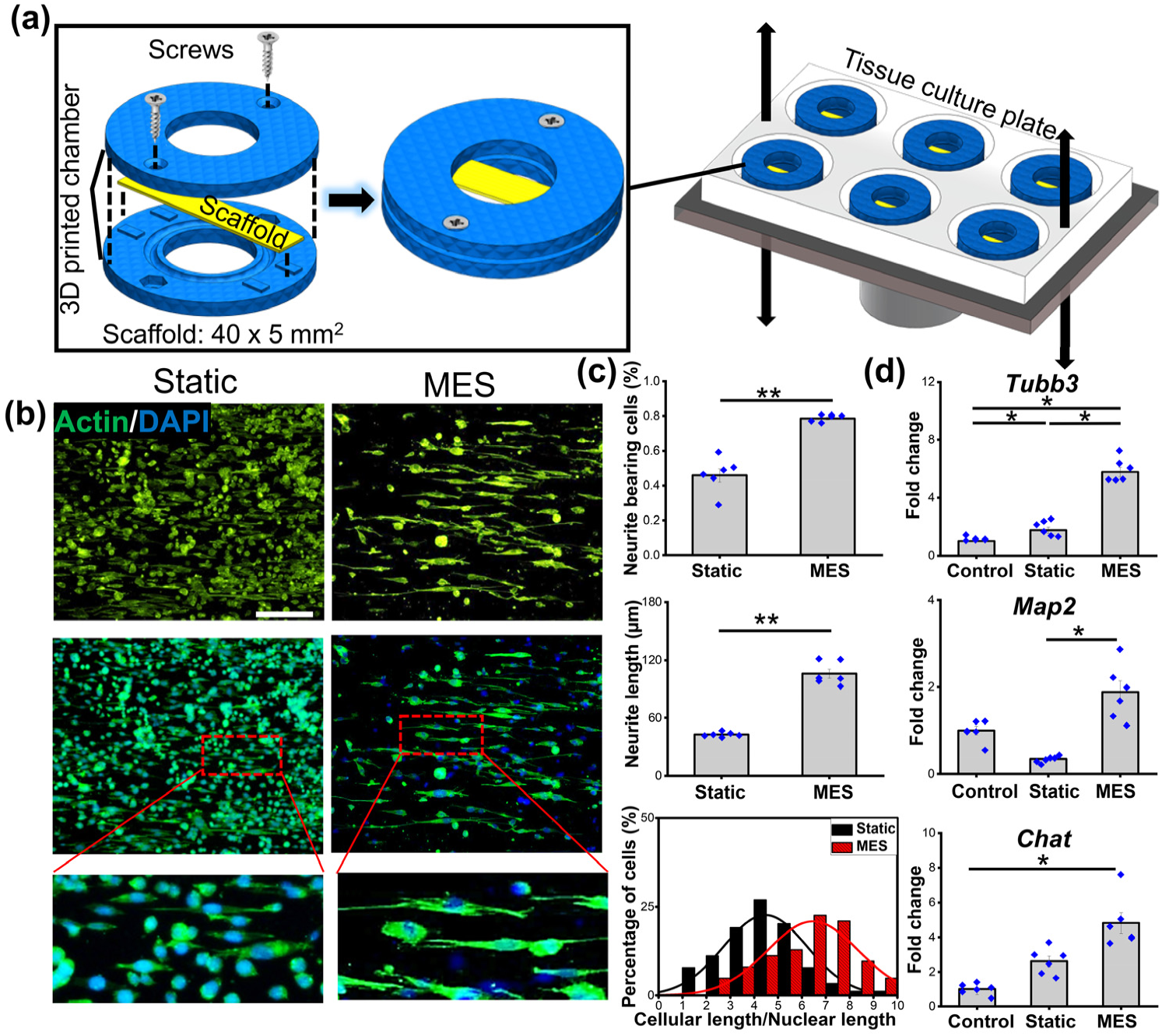
Effects of mechano-electrical stimulation (MES) on neurite formation, extension and neuronal maturation. (a) Schematic of MES setup describing the assembly of the hollow-cylindrical chamber that enables the deflection of the electrospun P(VDF-TrFE) scaffolds clamped in between under the hydro-acoustic actuation. (b) Representative immunofluorescence images showing cellular morphology of PC12 cells subjected to a 7-day culture with MES (2 hours per day). Cells cultured on P(VDF-TrFE) scaffolds without MES were used as the static control group (scale bar: 100 µm). (c) The percentage of cell population bearing neurites, average neurite length, and the percentage of neurite-bearing cells possessing a certain range of cellular length-to-nuclei length ratio, quantified from immunofluorescence images in (b) (* and ** denote statistical significance of p < 0.05 and p < 0.01, respectively. Six fluorescence images from independent samples were used for each condition). (d) The relative gene expression levels of neuronal markers (*Tubb3*, *Map2*, and *Chat*) under static or MES condition. The gene expression level was normalized to that of the cells that were cultured on tissue culture plates (Control) (n=6, * denotes statistical significance of p < 0.05).

We further examined the effect of MES on Schwann cells, the other major cellular component in the peripheral nervous system besides neurons. RSC96 cells, a rat Schwann cell line, cultured statically on the electrospun P(VDF-TrFE) scaffolds grew in colonies, similar to those cultured on tissue culture plates. In contrast, a single-cell formation was observed after cells were subjected to MES (**Figure 4a**). In addition, the expression of nerve growth factor (NGF) was significantly greater under MES as compared to the Static condition (**Figure 4a, b**). Consistent with the immunofluorescence imaging results, the gene expression of *Ngf* was significantly upregulated by MES (**Figure 4c**). The application of MES also induced significant upregulation in the gene expression of mature and myelinating Schwann cell markers, *Krox20*, and *Pmp22* while there was no significant change of an early immature Schwann cell marker expression *Ncam-1* (**Figure 4c**).

**Figure 4.**
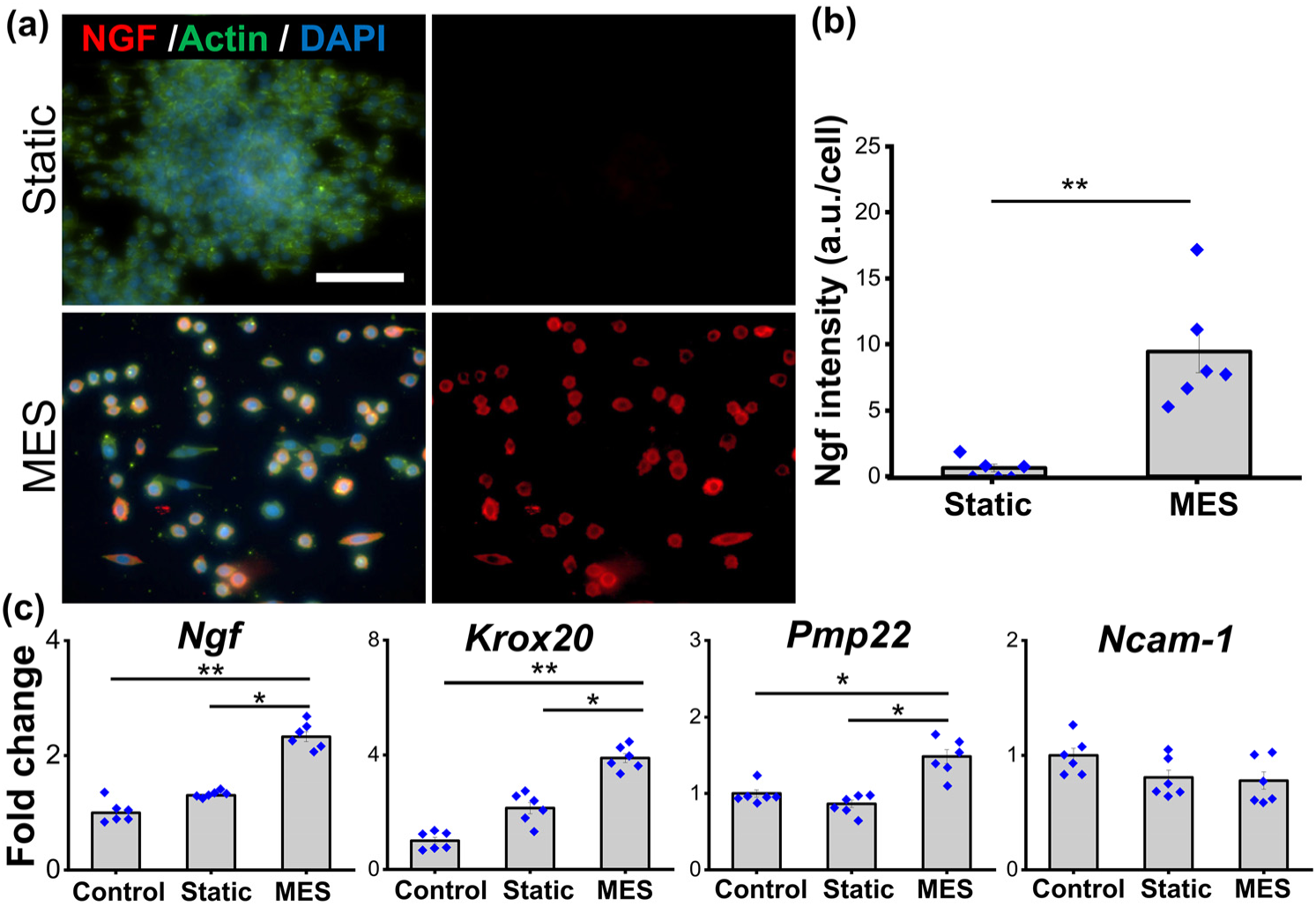
Effects of mechano-electrical stimulation (MES) on Schwann cell behaviors, including NGF secretion and cell maturation. (a) Immunofluorescence images showing NGF expression of RSC96 cells, a Schwann cell line, cultured statically or subjected to MES on P(VDF-TrFE) scaffolds (scale bar: 100 µm) and (b) their quantification of the fluorescent intensities (Six fluorescence images from independent samples were used for each condition). (c) The relative gene expression of neurotrophic factor (*Ngf*), myelination markers (*Krox20*, *Pmp22*), and early Schwann cell marker *Ncam-*1 in RSC96 subjected to MES for 7 days as compared to that of the statically cultured cells (n = 6, * and ** denote statistical significance of p < 0.05 and p < 0.01, respectively).

### 2.4. The effects of mechano-electrical stimulation on peripheral nerve regeneration in vivo

Based on the pro-neuronal and pro-glial effects of MES in vitro, we evaluated whether the physical stimulation by using the piezoelectric scaffolds will promote functional nerve regeneration in a rat model of peripheral nerve injury with a critical-sized defect. In order to accurately control the magnitudes of MES in vivo, the electrical outputs from P(VDF-TrFE) conduits, synthesized by rolling a P(VDF-TrFE) mat to form a hollow cylinder, were measured. The piezoelectric conduit with gold sputter-coated electrodes connected by insulated wires was implanted at the anatomical location near a sciatic nerve in a rat cadaver. To activate the piezoelectric conduit in a physiologically safe manner, an extracorporeal shockwave system, often used for the therapeutic management of musculoskeletal pains, was utilized. As expected, the voltage outputs from P(VDF-TrFE) conduits were directly proportional to the applied pressure magnitude of the shockwave stimulation, where a voltage output of 100 mV_p-p_ under 3 bars of shockwave increased to 400 mV_p-p_ under 5 bars (**Figure 5a**). The shockwave pressure of 4 bars was used for our in vivo study to produce 200 mV_p-p_, which was determined to be optimal for neural cell behaviors in the previous in vitro experiments (**Figure 5b**). A rat sciatic nerve transection model, proximal to the bifurcation of tibial and sural nerves was utilized. P(VDF-TrFE) conduit was implanted to bridge the transected nerves in a total of 8 rats, where they were subjected to either no additional treatments or periodical piezoelectric activation by shockwave application (**Figure 5c**). The rats were continuously monitored for peripheral nerve regeneration via walking track analysis (**Figure 5d**). Optical coherence tomography and histology were conducted as end-point analyses. Hereinafter, the 4 rats without further treatment (Static condition) will be referred to S1, S2, S3, and S4 while the 4 rats subjected to periodic shockwave treatment (MES condition) to MES1, MES2, MES3, and MES4.

**Figure 5.**
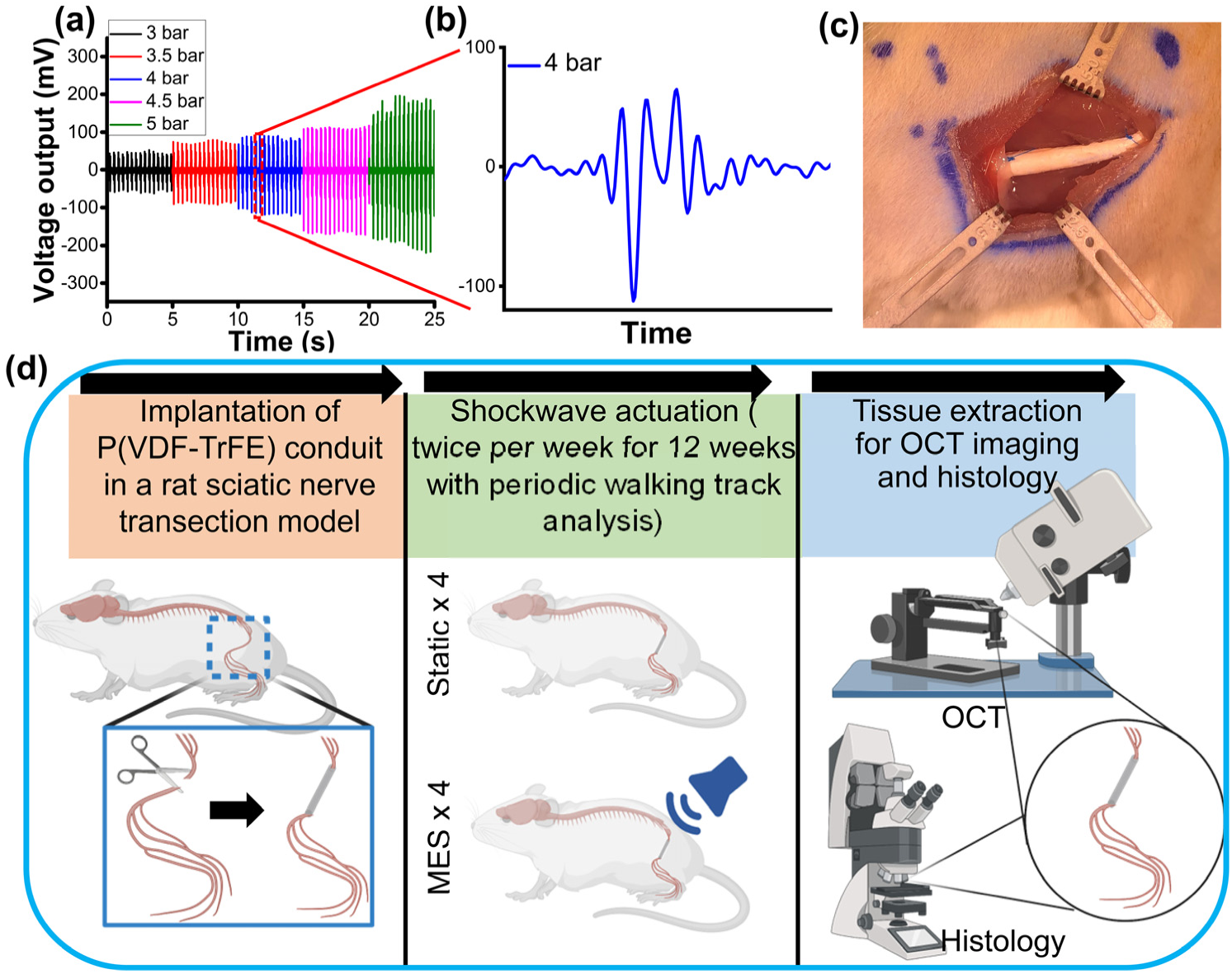
In vivo piezoelectric characterization of P(VDF-TrFE) conduits and the outline of animal experiments. (a) Shockwave magnitude-dependent voltage outputs from P(VDF-TrFE) conduits. (b) A zoomed-in voltage output graph showing the generation of 200 mV_p-p_ under the 4-bar pressure of the shockwave actuation. (c) A photograph showing the implantation of the P(VDF-TrFE) conduit into the rat to bridge the sciatic nerve gap. (d) A schematic describing the workflow of animal experiments to determine the efficacy of P(VDF-TrFE) conduit-mediated mechano-electrical stimulation on peripheral regeneration in a rat sciatic nerve transection model.

During the course of this animal study, we assessed the locomotive recovery in the rats by walking track analysis. There was a substantial improvement in toe spreading under the MES condition, as compared to that under the Static condition (**Figure S2a, Video S1, and Video S2**). The sciatic function index (SFI) with a range of 0 (healthy) and −120 (non-functional) was utilized to quantify the degree of functional regeneration at day 20, day 40, and day 80 post-surgery. A significant increase of the SFI value was observed from day 20 to day 80 under the MES condition while the Static condition showed no significant difference among the three time points (**Figure S2b**), indicating better functional regeneration by MES. It should be noted that the SFI values of Rat S3 and S4 (average of approximately −75) were higher than −120, the normalized value right after the sciatic nerve transection injury, indicating that the static control group exhibited a slight nerve function recovery probably due to the therapeutic effects of P(VDF-TrFE) conduits by themselves as demonstrated by others^27^. Nevertheless, the application of MES by the shockwave-activation of piezoelectric conduits resulted in a 150% improvement over the simple structural bridging.

To closely examine the MES-derived nerve regeneration, the whole nerve tissue/conduit samples were excised from the animals at the end of the study for further analyses. No visible inflammation was observed in either the Static or MES conditions (**Figure 6a, b**). In the longitudinal cross-sections, all rats subjected to MES (MES1-4) showed tissue connections while only 1 out of 4 rats exhibited a tissue connection within the conduit in the Static condition (S1-4). Even with this connection, however, axons (NF200, red color) were sparsely observed in Rat S2, as compared to relatively stronger axon staining intensities in MES1 - MES4 (**Figure 6c, d**). Moreover, the alignment of the regenerated and connected axons was readily observed in Rats MES1 - MES4, similar to healthy control (**Figure 6e**). To assess the detailed tissue structure of the regenerated nerves including axonal extension and myelination, confocal imaging was utilized (**Figure 6f, g, h**). In the Static group, axons and their myelination were sparsely observed with a disorderly organization, especially at the locations in the middle of the conduit and near the distal end except in Rat S2, where localized axonal alignment and myelin sheath co-expression was detected (**Figure 6f**). On the contrary, there was a considerably greater degree of aligned axon-myelin sheath formation spanning from the proximal end to the distal end in the MES group, comparable to the structure observed in the healthy control group (**Figure 6g, h**). It should be noted that the weaved structure observed in the healthy control group is likely due to the loss of tension during sciatic nerve extraction from the rat body, causing the coiling of the structure (**Figure 6e, h**). Taken together, the robust axonal extension for the proximal-distal re-connection and Schwann cell myelination was achieved by the application of MES.

**Figure 6.**
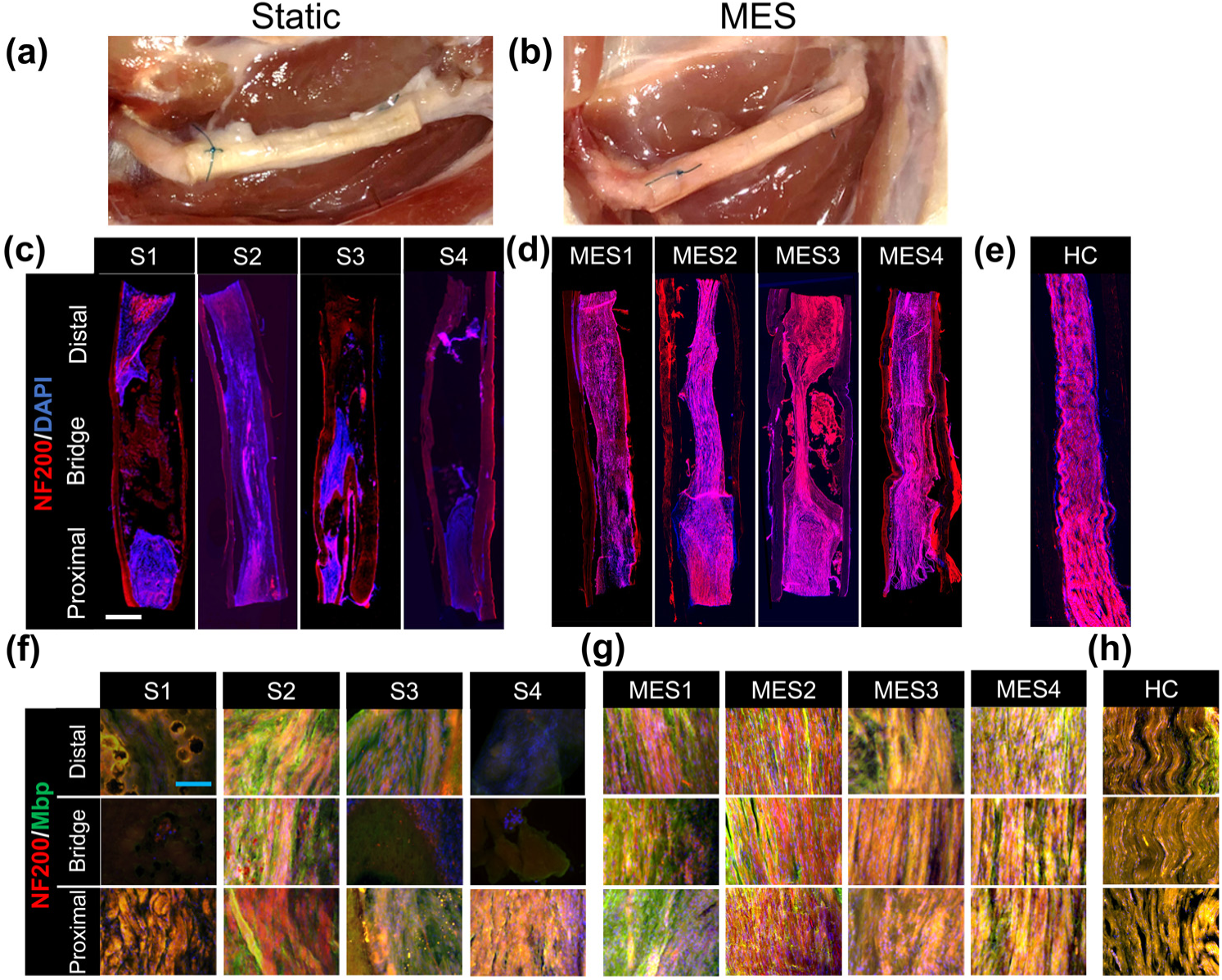
Morphological characterization of sciatic nerve regeneration by nerve re-connection. (a, b) Macroscopic observation of the P(VDF-TrFE) conduit 12 weeks post-surgery in (a) static and (b) mechano-electrical stimulation (MES) conditions. (c, d) Large-field-of-view immunofluorescence images showing the entire structure of P(VDF-TrFE) conduit and ingrowth tissue, bridging transected sciatic nerve in (c) Static and (d) MES conditions. The conduit/tissue samples were stained with an axonal marker NF200 and counterstained with DAPI. S1-S4 denotes each of the 4 rats in the Static group while MES1-MES4 denotes each of the 4 rats from the MES group (scale bar: 1 mm). (e) Morphology of a healthy sciatic nerve (healthy control (HC)). (f, g) High magnification images showing the tissue structure of the regenerated nerves at the proximal end, middle (bridge), and the distal end of the P(VDF-TrFE) conduits bridging transected sciatic nerves in (f) Static and (g) MES conditions. The conduit/tissue samples were double stained with an axonal marker NFH and a myelination marker MBP (scale bar: 100 µm). (h) Representative high-magnification images from the HC group.

We then evaluated the detailed myelination structure, a marker for functional recovery, by imaging the sciatic nerves in the transverse cross-sections at the proximal and distal ends of the nerve tissue/conduit samples. A significant number of damaged ovoid, composed of degraded axons and myelin sheaths^28, 29^, was observed in the Static group, as compared to the healthy control and MES condition (**Figure 7a, b Figure S3a, b**). Both the healthy control group and MES group showed significantly greater numbers of myelinated nerve fibers at the proximal and distal ends as compared to the Static condition (**Figure 7c**). The average axon diameter of the rats in the MES group was statistically larger than that in the Static group, while the healthy control group possessed the largest axon diameters (**Figure 7d**). Interestingly, the nerve structure at the proximal end in the MES group exhibited more uniform distribution of myelination, with better structural integrity, as compared to the tissue structure at the distal end, potentially indicating that a longer treatment may further improve nerve regeneration. Corroborating with the results from the bright field images, scanning electron microscopy (SEM) also showed a minimal number of normal healthy axons in the Static group while the morphology of the nerve in the MES group, especially at the proximal end, was highly comparable to that in the healthy control (**Figure S4**).

**Figure 7.**
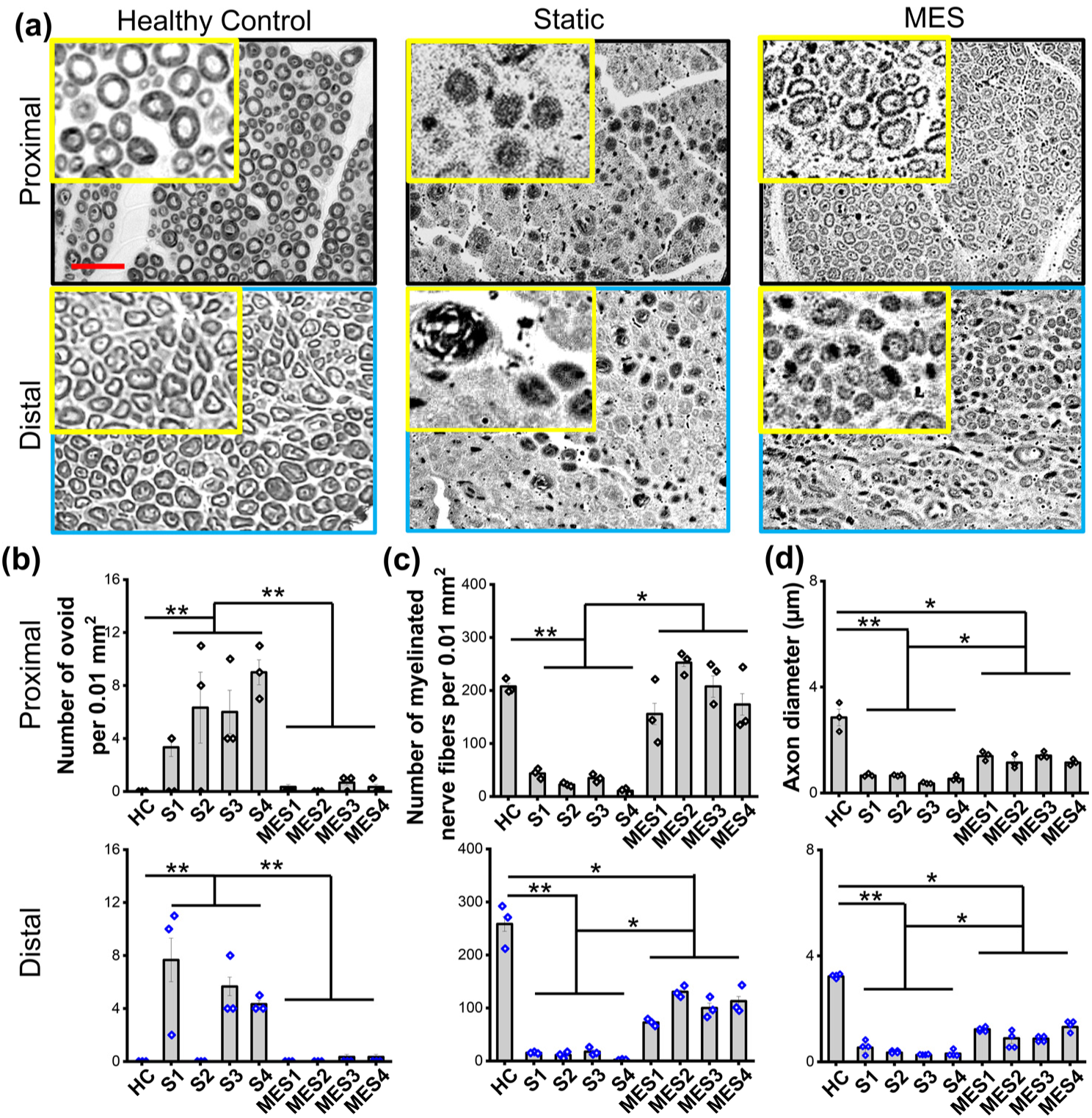
Morphological characterization of sciatic nerve regeneration. (a) Representative bright field images of axons and myelin sheaths at both the proximal and distal ends of the sciatic nerves within P(VDF-TrFE) conduits without (Static) or with shockwave actuation (MES). Cross-sectional images of sciatic nerves from healthy rats were used as the healthy control (scale bar: 20 µm). Quantification of (b) the number of myelinated nerve fibers, (c) the number of ovoid, and (d) axon diameter from histology images in (a). At least 3 images from each rat were used for the quantifications (* and ** denote statistical significance of p < 0.05 and p < 0.01, respectively).

### 2.5. Characterization of peripheral nerve regeneration by polarization-sensitive optical coherence tomography (PS-OCT) imaging

Polarization-sensitive optical coherence tomography (PS-OCT) was also used to non-invasively examine nerve repair and quantitatively assess the regenerative outcomes under Static and MES conditions. Nerve exhibits birefringence30, an optical property of materials, when polarized light passes through a birefringent material and its two constituent orthogonal polarization components travel at different speeds due to the difference of their refractive indices, resulting in phase retardation. PS-OCT measures phase retardation and uses this measurement to generate image contrast^31, 32^. In particular, due to the presence of anisotropic myelin sheath, the nerve exhibits form birefringence33, which arises from fibrous materials consisting of long, parallel fibrils embedded in a medium of different refractive index. As a consequence, a change in the density and organization of the myelinated nerve fibers would result in a change in birefringence which can be detected using PS-OCT^34–36^. We scanned the nerve-conduit samples using a spectral-domain PS-OCT system and generated intensity and phase-retardation images using the Stokes vector-based method^37, 38^. Representative 3D reconstructed PS-OCT images of the Static and MES conditions are presented in **Figure 8a, b**. In these images, the structural intensity of the conduit is presented in semi-transparent grayscale and the phase retardation of the nerve is presented in color. In the intensity images, whiter colors represent higher back-reflected light intensity whereas in the phase retardation images, blue to red color indicates phase retardation in the range [0.07 0.2] deg µm^−1^. As evident in the images, full nerve connectivity from the proximal (left) to the distal (right) end was observed from the high-phase retardation regions colored in yellow and red throughout the length of the sample under the MES condition. On the other hand, the transected gap is still present between the proximal and distal ends within the conduit under the Static condition. These images are in agreement with the longitudinal histological sections in **Figure 6**, but present volumetric 3D images of the nerve inside the conduit for the whole length of the samples, demonstrating the significant potential of PS-OCT imaging for non-destructive monitoring of nerve repair.

**Figure 8.**
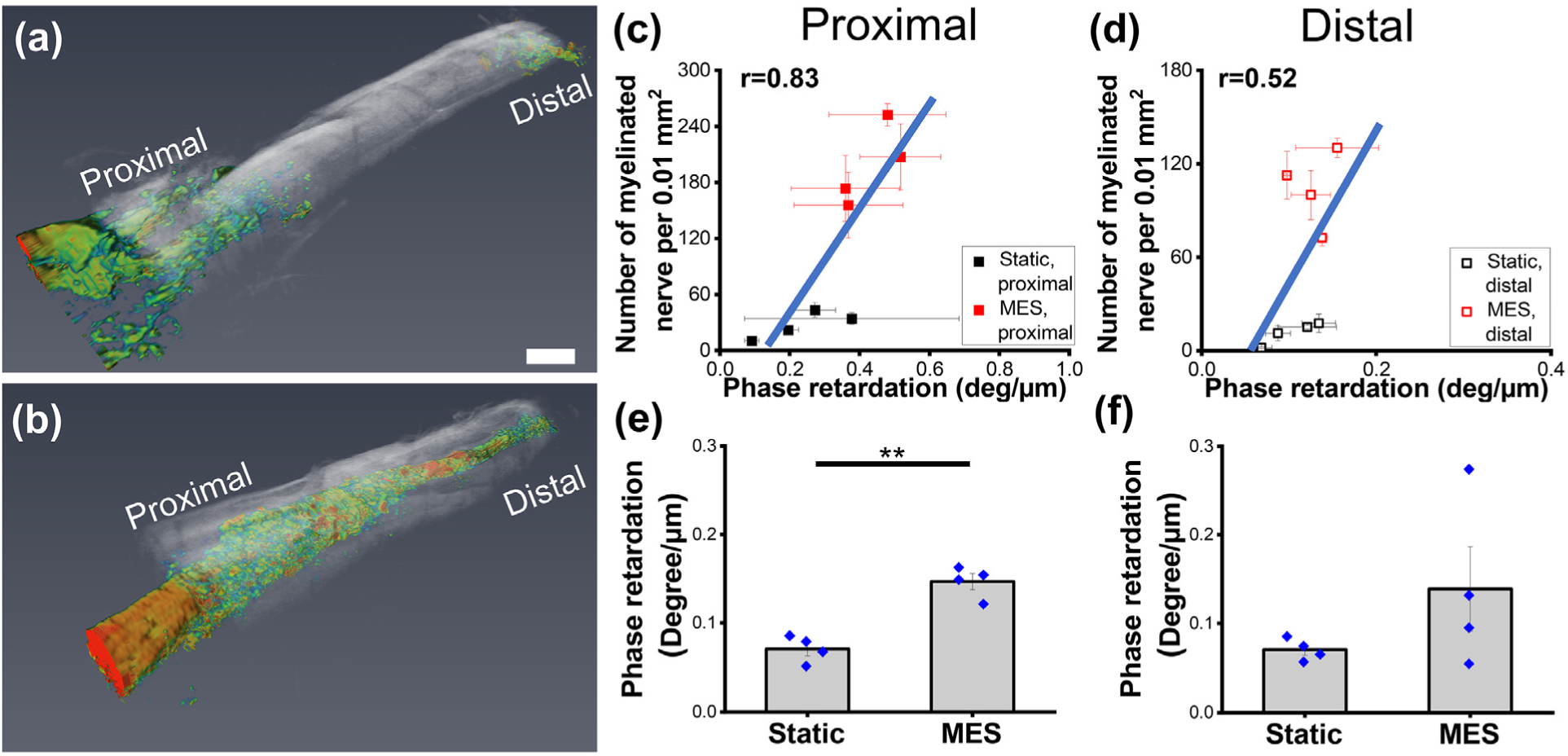
Polarization-sensitive optical coherence tomography (PS-OCT) imaging of the sciatic nerves. (a, b) Representative PS-OCT images after 3D reconstruction of the sciatic nerves without or with MES. The structural intensity image in grayscale shows the conduit surface whereas the colored phase retardation image shows the nerve. In the intensity images, whiter colors represent higher back-reflected light intensity. In the phase retardation images, blue to red color indicate low to high phase retardation in the range [0.07 0.2] deg µm^−1^ (Scale bar: 2 mm). (c, d) Correlation between the number of myelinated nerve fibers per 0.01 mm^2^, quantified from Figure 7, and the phase retardation of the nerve outside of the conduit in the proximal and distal ends, respectively. (e, f) Quantification of the phase retardation of the nerve/tissue inside of the conduit at the proximal and the distal ends, respectively (** denotes the statistical significance of p < 0.01).

For quantitative comparison, the average phase retardation slope, which is proportional to birefringence^39^, was measured. Birefringence is a measure of the degree of myelination of the nerve which is evident from the PS-OCT study of nerve crush injury in rats where a decrease in myelination of the nerve was shown to result in a decrease in birefringence^35^. We measured the average phase retardation slope from tiles of 340 × 340 x 2 µm^3^ regions from both the proximal and distal ends outside of the conduit for each sample. These values were plotted (**Figure 8c, d**) against the number of myelinated nerve fibers per 0.01 mm^2^, which is a measure of nerve functionality as described above in the histological results obtained from the transverse cross-section. From these plots, we observed a positive correlation between the two parameters in both the proximal and the distal ends (r = 0.83 and 0.52, respectively) consistent with the expectation that an increased number of myelinated nerve fibers would result in increased birefringence.

When comparing the phase retardation slope of the nerve inside the conduit for the different rats using a similar measuring criterion (**Figure 8e, f**), we observed a significantly higher (p = 0.0006) phase retardation slope in the rats under the MES condition compared to the Static ones at the proximal end. This indicates that the myelinated nerve fibers are denser and more organized at the proximal ends in the rats under MES condition which is in agreement with the histological images in **Figure 6**. The distal end also showed an appreciable increase in the phase retardation value in the MES condition.

## 3. Discussion

The treatment of critical-sized peripheral nerve injuries is uniquely challenging due to the complex nature of the nerve tissue, often resulting in limited functional recovery. Autologous nerve transplantation has been considered the gold standard for peripheral nerve treatment^9^. However, this method has several disadvantages, such as the necessity for multiple surgeries, the creation of a functionally impaired region where the graft was taken from, the disproportion of the graft to nerve tissue in size and structure, and most substantially, the high risk of neuroma formation^40^. As an alternative approach, various types of natural biomaterials, including collagen, chitosan, gelatin, and silk fibroin, are used as nerve conduits to bridge the injured nerve gaps, leveraging their excellent biocompatibility and biodegradability. However, the poor mechanical properties with a typically rapid degradation rate, limit the use of natural materials as nerve guidance conduits in vivo^3, 41^. To overcome this limitation, numerous biocompatible synthetic materials were developed as nerve guidance conduits with certain physical properties to achieve the desired peripheral nerve regeneration^42–44^. These synthetic conduits provide additional functionality to the structural support of the conduits to further enhance nerve regeneration. Several studies have attempted to use electrical stimulation using conductive materials as nerve guidance conduits to promote peripheral nerve regeneration^22^. A major drawback of this approach, however, is the use of external electrical wires connecting the conduit inside of the body and the power source outside, causing technical complexity and significant risks for complications including infection^22, 45^. A recent novel study has demonstrated the use of radio frequency-activated electronic devices for the non-invasive electrical stimulation of nerves in vivo^46^. In spite of its positive phenomenological outcomes, a short in vivo life span due to biodegradation and likely limitation in the magnitudes of the electrical stimuli generatable from the device is uncertain for its translational applications.

In this regard, piezoelectric polymer, especially PVDF and its derivative P(VDF-TrFE), is a highly potent material for non-invasive electrical generators^23, 26^. There have been several attempts to utilize the piezoelectric effect for nerve regeneration^47^. An animal study demonstrated a certain degree of enhancement in nerve regeneration by electrically poled (to improve piezoelectricity) PVDF conduits^27^. More recently, a group of researchers exploited the enhanced piezoelectricity of P(VDF-TrFE) scaffolds synthesized by electrospinning to accelerate neurite outgrowth of dorsal root ganglion neurons or promote neuronal differentiation of human neural stem/progenitor cells in vitro^48, 49^. Notably, P(VDF-TrFE) conduits seeded with Schwann cells were utilized to bridge the spinal cord injury and showed enhanced axon regeneration as well as improved myelination^50, 51^. In spite of favorable results, these studies were likely unable to utilize the true potential of piezoelectricity as it requires dynamic straining of the materials to generate electric potentials. In this regard, we aimed to develop a methodology to activate piezoelectric P(VDF-TrFE) conduits by external mechanical actuations in a physiologically safe manner to efficiently and non-invasively apply the combination of electrical and mechanical stimulation for enhanced nerve regeneration in vivo.

This limitation in utilizing piezoelectric materials for electrically stimulating cells/tissues is partly due to the inefficiency of the materials in converting the applied mechanical actuation to electrical energies or low piezoelectric coefficients; a very high magnitude of mechanical strain, which would detrimentally affect cells/tissues, is necessary to generate physiologically meaningful magnitudes of electric potentials. We overcame this limitation by achieving transformative enhancement in the piezoelectric properties of electrospun P(VDF-TrFE) nanofibers through material/process optimization^23^. This improvement enables the generation of electric potentials in appropriate magnitudes to stimulate neural cells by physiologically safe mechanical actuation. Based on our recent finding that the magnitude of the piezoelectric potential generation of electrospun P(VDF-TrFE) can be modulated by controlling the fiber diameter and scaffold thickness as well as the magnitude of applied strains^26^, a piezoelectric conduit composed of P(VDF-TrFE) fibers was optimized to produce physiologically relevant magnitudes of electric potentials. Since the size of the nanofibers affects cellular alignment and subsequent neurite extension the fiber diameter was carefully optimized by balancing the fiber diameter-dependent cellular behaviors and piezoelectric properties.

The potential of piezoelectric scaffold-mediated mechano-electrical stimulation for promoting functional neural cell development was tested in vitro. The results demonstrate that mechano-electrical stimulation not only significantly improves neuronal differentiation and neurite outgrowth but also induces the maturation of Schwann cells with enhanced secretion of NGF, a potent neurotrophic factor. These results are consistent with previous studies showing the regenerative effects of electrical stimulation and mechanical stimulation in neuronal cells and glial cells, respectively^21, 52^. To activate the piezoelectric effect of electrospun P(VDF-TrFE) conduits in vivo, a therapeutic shockwave system was utilized to apply mechanical actuation to the implanted conduits. Therapeutic shockwave generates acoustic waves with an energy density between 0.01 and 0.6 mJ mm^−2^ and a penetration depth of over 40 mm. It has been therapeutically used for musculoskeletal pain treatment in animals and humans^53^. The use of acoustic waves for the activation of the piezoelectric conduit removes the need for invasive electrodes for the functional regulation of neural cells. The magnitude of the shockwave magnitude was adjusted to induce the optimized voltage output. The application of mechano-electrical stimulation during the 12-week in vivo study resulted in significant recovery in motor function, observed by walking track analysis. According to histology on the longitudinal nerve sections, mechano-electrical stimulation substantially enhanced the reconnection of the severed sciatic nerve (4 out of 4 rats), as compared to the control Static group where the sciatic nerve gaps were bridged with P(VDF-TrFE) conduits but without the application of mechano-electrical stimulation (1 out of 4 rats). The sciatic nerve cross-sections in the transverse direction also showed that there was a superior axon and myelin regeneration by MES. Although there are other studies that showed peripheral nerve regeneration by various techniques, the maximum defect gap size was usually limited to 10 mm or smaller, often resulting in poor end-to-end connection with a lack of functional regeneration^13, 54^. In contrast, we showed the capability of mechano-electrical stimulation, fully connecting the transected sciatic nerve with a larger critical-sized defect in rats. Combined with the results from morphological and functional analyses, the superior clinical potential of piezoelectric conduit-mediated nerve regeneration is demonstrated.

PS-OCT imaging technique was further used to confirm peripheral nerve regeneration. By measuring the phase retardation arising from the birefringence in the nerve, PS-OCT images, and subsequent analysis proved the existence of full nerve connection under the MES condition while very poor or no connections were observed under the Static condition. More interestingly, there was a close correlation between the number of myelinated axons from the histological cross-sectional images and the phase retardation slope from the PS-OCT images. In addition, the phase retardation values at the proximal end inside the conduit showed a significant difference between the MES and Static conditions. The range of phase retardation values in the samples here is consistent with that found in previous studies^34, 35^, where PS-OCT was used to quantitatively assess the condition of injured and healthy sciatic nerves in rats. Although the phase retardation values of our nerve samples are on the lower side compared to the healthy sciatic nerves^34^, the values in the MES group align with week 3 and 4 post-injury sciatic nerves^35^. Since PS-OCT non-invasively acquires 3D volumetric images, it provides the added benefit of virtually slicing the sample in any plane of interest for qualitative and quantitative analysis. Considering all, this demonstrates the ability and versatility of using the PS-OCT technique for morphological and functional assessment of the sciatic nerves without any potential artifacts and sampling errors that usually accompany tissue sectioning.

## 4. Conclusion

In this study, we developed and validated a method to non-invasively apply mechano-electrical stimulation to the injured sciatic nerves for their enhanced regeneration by utilizing the long-term in vivo activation of piezoelectric conduits. After the scaffold optimization of physical characteristics based on neuronal differentiation, neurite outgrowth, and Schwann cell maturation in vitro, we further found that the subjection of mechano-electrical stimulation, mediated by the periodic activation of piezoelectric conduits, significantly promoted axonal regeneration, myelin regeneration, and functional motion recovery in vivo. Collectively, given its non-invasive physical stimulation and straightforward implementation, the mechano-electrical stimulation system presented in this study has significant potential for clinical translation, potentially offering an efficient therapeutic solution for peripheral nerve injuries.

## 5. Experimental Section

### Synthesis of electrospun P(VDF-TrFE) scaffolds

P(VDF-TrFE) scaffolds with various fiber diameters were electrospun as described elsewhere (ref). Briefly, A solution of 5, 7, or 16 wt. % P(VDF-TrFE) (70:30 mol%, Solvay), fully dissolved in a solvent system of 60/40 volume ratio of N,N-dimethylformamide (DMF) (Sigma) to acetone (Fisher), supplemented with 1.5 wt. % of pyridinium formate (PF) buffer (Sigma), was electrospun to produce fibers having an average fiber diameter of 200, 500, or 800 nm, respectively. Various parameters, including needle tip to collector distance (approximately 10 cm), applied voltage (−15 kV to −20 kV), solution feed rate (6 mL h^−1^), and absolute humidity (7.6 g cm^−3^) at room temperature (23 °C), were optimized for a stable and consistent electrospinning process. A rotating wheel with an angular velocity of 47.9 m s^−1^ was used to collect the electrospun fibers to yield an aligned fibrous structure. The electrospinning duration was adjusted for each polymer concentration to produce scaffolds with approximately 200 µm thickness. The electrospun P(VDF-TrFE) was subsequently annealed at 90 °C for 24 h to further enhance its piezoelectric properties^23^.

### Piezoelectric characterization of electrospun P(VDF-TrFE)

The d_33_ piezoelectric coefficient was determined for P(VDF-TrFE) fibers with various diameters following a previously described method^26^. Briefly, nanofibers of P(VDF-TrFE) were sparsely deposited on a thermal-oxide silicon substrate coated with gold and subjected to the single-point piezoresponse force measuring mode using an MFP-3D atomic force microscope (AFM) (Asylum research). Multiple points were analyzed on each scanned fiber and the piezoresponse (voltage and amplitude) measurements were taken by applying a voltage bias between the tip (AC240TM, Asylum) and the collector. The value of the d_33_ was calculated by,

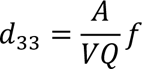

where A is the piezoresponse amplitude, Q is the quality factor influenced by the currently used AFM probe, and f is the correction factor determined by a standard piezoelectric sample with a known d_33_ value.

To quantify the voltage outputs from the electrospun P(VDF-TrFE) scaffolds, a different magnitude of mechanical actuation was applied onto acellular scaffolds, having a dimension of 45 × 5 mm^2^, positioned inside the cell culture system as described in **Figure 3a**. The scaffolds were gold-coated on both sides for electrodes and connected to an oscilloscope (Pico Technologies) for real-time voltage signal measurement. The corresponding strain change was determined by our customized cantilever system^26^.

### Cell culture

PC 12 cells, a rat neuronal progenitor cell line (ATCC), were maintained in type I collagen (Sigma)-coated tissue culture plates with culture media composed of F12K (ATCC), 15% horse serum (Gibco), 2.5% fetal bovine serum (FBS, VWR), and 1% antimycotic/antibiotic solution (Corning). For neuronal differentiation, the media was changed to F12K, supplemented with 1% FBS, 50 ng mL^−1^ nerve growth factor (NGF), and 1% antimycotic/antibiotic. The cells were allowed to pre-differentiate for 48 hours in tissue culture plates before seeding them onto electrospun P(VDF-TrFE) scaffolds.

RSC96 cells, a rat Schwann cell line (ATCC), were cultured in DMEM (Corning), supplemented with 10% FBS and 1% penicillin/streptomycin (Meditech). The cells were cultured and maintained in tissue culture plates before their inoculation onto the scaffolds.

### The effect of scaffold morphology and electrical stimulation on PC12 cell behaviors

Pre-differentiated PC12 cells were seeded onto electrospun P(VDF-TrFE) scaffolds having fiber diameters of 200, 500, or 800 nm at a seeding density of 40,000 cells cm^−2^. The cells were cultured for 72 hours, before being fixed in 4% paraformaldehyde (PFA) for imaging analysis.

To determine the optimal electrical stimulation regimen for peripheral neurogenesis, pre-differentiated PC12 cells were seeded onto electrospun P(VDF-TrFE) scaffolds having the optimized fiber diameter (500 nm), and subjected to various magnitudes of single-day or multi-day electrical stimulation. The setup for electrical stimulation was previously described^25^. Briefly, the cell/scaffold construct was placed on a conductive surface (gold-coated polystyrene) within a well of the 12-well tissue culture plate (**Figure 2a**). The bottom of each well was drilled to create a hole, enabling electrical wires to connect between a function generator and the cell/scaffold construct. A grounded gold-coated glass coverslip was placed above the gold-coated polystyrene in contact with the cell culture media within each well. Various magnitudes of electrical signals (100, 200, and 400 mV_p-p_), with a signal shape similar to the voltage output peaks generated from the mechanical actuation-activated piezoelectric P(VDF-TrFE) scaffolds, were applied to the cells. The PC12 cells were either electrically stimulated once for 2 hours, followed by 70 hours of static culture, or daily stimulated (2 hours/day) for three days. For both conditions, the cells were cultured for a total duration of 72 hours before fixation with 4% PFA.

### The effect of mechano-electrical stimulation on PC12 and RSC96 cell behaviors

To demonstrate the feasibility of mechano-electrical stimulation in promoting peripheral nerve cell behaviors, the pre-differentiated PC12 cells or the Schwann cells were separately subjected to mechano-electrical stimulation. P(VDF-TrFE) scaffolds, inoculated with the cells, were subjected to hydroacoustic actuation, based on the optimal parameters determined above, to realize the piezoelectric effect (**Figure 3a**). The voltage outputs generated from the scaffolds were controlled to be approximately 200 mV_p-p_. The cells were daily stimulated for 2 hours for 7 consecutive days before fixation in 4% PFA or lysed for gene expression analysis.

### Fluorescence imaging

PC12 and RSC96 cell/scaffold constructs were stained with 4′,6-diamidino-2-phenylindole (DAPI, Sigma) and phalloidin-Alexa Fluor 594/488 (Abcam) to visualize cellular morphology. A fluorescence microscope (Nikon) was used to image the stained cells. For PC12 cells, the number of cells exhibiting neurites (having at least one neurite with a length equal to the cell body diameter^55^), average neurite length, and the distribution of neurite lengths to nuclear lengths ratio were measured using the ImageJ software. For RSC96 cells, the fluorescence intensity of neurotropic NGF expression was measured.

### Gene expression analysis

The effects of mechano-electrical stimulation on PC12 and RSC96 cells after 7 days of culture were determined by quantitative polymerase chain reaction (qPCR). A Rneasy Micro Kit (Qiagen, Valencia, CA) was used to isolate RNA from the lysed samples, followed by cDNA synthesis using an iScript cDNA Synthesis Kit (Bio-Rad, Hercules, CA). Real-time qPCR was performed to determine the expression levels of phenotypic markers (**Table S1**). Raw data were analyzed by the comparative threshold cycle (C_T_) using an endogenous control.

### Fabrication and electrical characterization of electrospun P(VDF-TrFE) conduits for in vivo experiments

After synthesizing electrospun aligned P(VDF-TrFE) scaffold as described above, the electrospun mat was rolled onto a metal rod to form a cylindrical conduit, with the aligned fibrous direction matching to the longitudinal direction of the conduit. A medical adhesive (Factor II) was used to glue the two long edges of the scaffold together to form a conduit with an inner diameter of 1.5 mm and a length of 17 mm, matching the size of the rat sciatic nerve with a critical injury gap size of 15 mm.

To determine the magnitude of shockwave for the generation of an electrical output of 200 mV_p-p_, the voltage outputs of the conduits implanted in the rat body were characterized. Briefly, both sides of the electrospun P(VDF-TrFE) scaffolds were sputter-coated with gold as electrodes before forming the conduits. The gold-coated conduit was implanted into a sacrificed rat at the peripheral nerve injury site, with two electrical wires connecting the inner and outer surfaces of the conduit to the oscilloscope. The voltage signal data was detected and recorded under various magnitudes of the shockwave application.

### Surgical procedures

In this study, all procedures were approved for animal experiments from UCR IACUC (animal use protocol 20210016). Adult Sprague Dawley rats (n = 8, Taconic) weighing approximately 300 g, were randomly divided into two groups. Rats were subjected to sciatic nerve transection surgery and the severed ends were sutured to the P(VDF-TrFE) conduit to form a nerve gap of 15 mm. 20 mg mL^−1^ Matrigel solution (Corning) was injected into the conduits after suturing to ensure a suitable biological environment for axon regeneration. Experimental rats (MES group, n = 4) were subjected to periodic shockwave applications while the control group (Static group, n = 4) was maintained without the application of mechano-electrical stimulation after the surgery. For the MES group, 1000 shockwave pulses, with a magnitude of 3.5 bar and a frequency of 3 Hz, were applied near the implanted conduit twice a week for a total of 12-week duration.

### Walking track analysis

On day 20, day 40, and day 80 post-surgery, the walking track analysis was conducted to assess the motion recovery of the rats^56^. From the snapshots of the footprints recorded from underneath the walking platform, several length measurements, including the width of the toe spread (TS) and print length (PL), were taken using the ImageJ software. The sciatic functional index was calculated based on these measurements using the equation^57, 58^ below,

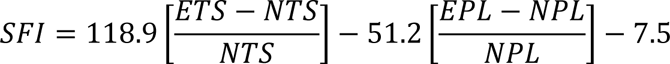

where E stands for the experiment footprint while N stands for the normal footprint. An SFI of 0 is normal and an SFI of −120 indicates complete impairment. Note that rats S1, S2, MES1, and MES2 lost their digits due to autotomy, preventing the tracking of SFI over the course of the animal experiment.

### Nerve conduit harvesting

At the 12-week time point post-surgery, all rats were euthanized, and the implanted conduits were excised. The uninjured counternatural sciatic nerves were used as healthy controls. All the samples were fixed in a 2% glutaraldehyde/2% paraformaldehyde solution at 4°C for 48 hours for future analysis.

### Polarization-sensitive optical coherence tomography (PS-OCT) imaging

All fixed tissue samples were scanned with a custom-built spectral domain PS-OCT system, having a center wavelength of 1310 nm and an FWHM bandwidth of 68 nm. The system has an axial and lateral resolution of 11 µm and 37 µm, respectively. The whole length of each sample was scanned volumetrically in five overlapping sections, with each section having a field of view of 4.5 mm × 4.5 mm and approximately 1 mm overlap between subsequent sections which were manually aligned when generating an image of the full length of the sample. The volumes comprised 200 cross-sectional images, each of which had 512 A-lines. The scanning was done at a rate of 4435 A-lines per second with the fast-scanning axis being perpendicular to the axis of the nerve conduit sample.

The volumetric data were processed with MATLAB to generate structural intensity and phase retardation cross-sectional images. First, structural intensity images were generated following the standard Fourier domain processing method^30, 59–62^. From the intensity images, we manually traced the surface of the nerve in every ten frames in a volume which gave us the depth of the nerve surface at different A-lines in those frames. The depth values at different A-lines across every ten frames were then fit with a low-order polynomial to determine the nerve surface in every frame (**Figure S5**). These surfaces were used as the polarization reference to generate cumulative phase retardation images using the Stokes vector method^31^.

The height of the cumulative phase retardation curve was adjusted so that the nerve surface remains at depth 0. From the height-adjusted cumulative phase retardation vs depth plots, the slope of the rising portion of the curve, which is proportional to birefringence^39^, was measured. First, to reduce the effect of speckle, five (corresponds with the lateral resolution) neighboring height-adjusted cumulative phase retardation curves in the first scanning direction were averaged. Also, for reliable measurements, only regions with a high (> 0.5) degree of polarization uniformity (DOPU) were considered^63^. From the high DOPU regions, we have manually identified the endpoints of the rising portion of the height-adjusted cumulative phase retardation curves from the central A-line in the nerve/tissue region in every ten frames in a volume and used a low-order polynomial fit to find the endpoints in every frame (**Figure S6**). A linear least square fit was then used between these endpoints for calculating the slope.

The measured phase retardation slope values were used for quantitative comparison. For comparison with histological parameters, we first divided the nerve regions at the proximal and distal ends outside of the conduit into 340 × 340 × 2 µm^3^ tiles (**Figure S7**). From each tile, the average of the phase retardation slope values was measured. The mean and standard error of these measurements across the tiles were calculated for the proximal and distal ends separately for each rat and compared with histological measurements. Similar measurements were also taken from the nerve regions inside the conduit. For this, we have selected the second volume section from both the proximal and distal ends divided the nerve regions in these sections into 340 × 340 × 2 µm^3^ tiles and averaged the phase retardation slope values inside each tile (**Figure S7**). The mean and standard error of these measurements across the tiles were calculated for both the proximal and distal ends of the different rats and were used for the comparison between the Static and MES groups.

For visualization of the samples, we have used structural intensity and depth-resolved phase retardation images. To generate the depth-resolved phase retardation images, the slope of the cumulative phase retardation curve was measured locally for every depth pixel using the least square fitting. In this case, five neighboring phase retardation curves in the fast-scanning direction were averaged to reduce the speckle, and only regions with high (> 0.5) DOPU were considered for reliable measurements. The cross-sectional intensity and depth-resolved phase retardation images from the five overlapping sections were compiled in Amira 3D visualizing software for volumetric 3D visualization. The five sections were manually aligned for generating an image of the whole length of a sample. The intensity image was presented in grayscale and the phase retardation image was represented in color. In the intensity images, whiter colors represent higher back-reflected light intensity whereas in the phase retardation images, blue to red color indicates low to high phase retardation in the range [0.07 0.2] deg µm^−1^. For easy visibility of the phase retardation of the nerve inside the conduit, the intensity image was made semi-transparent.

### Histology

After the PS-OCT imaging, each conduit sample was cut into 3 pieces using a sharp scalpel, including an 11 mm length of the middle portion, a 3 mm length of the proximal portion, and a 3 mm length of the distal portion. The middle portion of the conduit was prepared for cutting sections in the longitudinal direction. Briefly, the samples of the middle portion were sequentially dehydrated in 15%, and 30% sucrose (Fisher), for 15 minutes each, followed by overnight incubation in optimal cutting temperature (OCT, Fisher) compound. The samples were then embedded in a fresh OCT compound and positioned in a way to enable longitudinal directional sectioning. The sample was snap-frozen in isopentane (Sigma) in a liquid nitrogen bath. The frozen block was then cut into longitudinal sections having 20 µm thickness using a cryostat (Leica). The sections were stored in the fridge until further histological analysis. For the immunohistochemical assessment of in vivo nerve regeneration, the longitudinal nerve-conduit sections were double-stained with neuronal marker NF200 (DSHB), myelination marker MBP (Bio-Rad), and counter-stained with DAPI. The samples were imaged under a fluorescence microscope with a large field imaging function to reveal the whole length of the section. The detailed neuronal and myelination structure was further visualized using confocal microscopy (Zeiss).

The proximal and distal portions of the conduits were used for the cross-sectional cut in the transverse direction of the nerve. Briefly, fixed samples underwent post fixation and staining in 2% osmium tetroxide dissolved in PBS, followed by a sequential dehydration process in 30%, 50%, 70%, and 100% ethanol (Fisher). The dehydrated samples were then incubated in a transitional solvent of 100% propylene oxide (Sigma) overnight and transferred to the Epon-812 compound (Electron Microscopy Sciences). The mixture was incubated in the oven at 60°C overnight to facilitate resin cross-linking. The hardened resin blocks were mounted onto an ultramicrotome (RMC) and cut into 1 µm thickness sections. The sections were stored in a fridge in preparation for bright field microscopy (Olympus). The number of intact axons with myelin sheaths, the number of ovoid formations, and the average axon diameter was measured and compared among the healthy control, Static, and MES groups.

### Statistical analysis

All experiments were conducted with a minimum of triplicate biological samples and data are presented as mean ± standard deviation (SD) or standard error of means, depending on the type of data obtained. Comparison of experimental groups for statistical significance was determined using the IBM SPSS software with either one-way ANOVA with Tukey’s HSD posthoc test or a two-sample student T-test. Statistical significance was reported when a ‘p’ value was less than 0.05.

## Supplementary information

Supplementary information (TablesS1, Figure S1-Figure S7)

Video S1. Representative rat walking video from the MES group at day 80 post surgery. Video S2. Representative rat walking video from the Static group at day 80 post surgery.

## Author contributions

YT performed in vitro and in vivo experiments, data analysis and wrote the manuscript. TIT performed OCT imaging and its analysis. SW and NB analyzed histology images. BHP and JN conceptualized the study and wrote/revised the manuscript.

## Acknowledgements

This study was partially funded by the National Science Foundation (CBET-1805975). The funder played no role in study design, data collection, analysis and interpretation of data, or the writing of this manuscript.

## Data availability

The datasets used and/or analysed during the current study available from the corresponding author on reasonable request.

