## Supplementary Information for "Enhanced Peripheral Nerve Regeneration by Mechano-electrical Stimulation"

**Table S1. Primer sets for rt-qPCR analysis of PC12 and RSC96 cells.**

| Marker | Primer | Forward | Reverse |
|---|---|---|---|
| Housekeeping | *Rps18* | 5'-CCCGAGAAGTTTCAGCACATC-3' | 5'-ATGGCAGTGATAGCGAAGGCT-3' |
| | *Gapdh* | 5'-GCAAGAGAGAGGCCCTCAG-3' | 5'-TGTGAGGGAGATGCTCAGTG-3' |
| Neuronal markers | *Tubb3* | 5'-GGGCCAAGTTCTGGGAAGTC-3' | 5'-AGTCGCCCACGTAGTTGCC-3' |
| | *Map2* | 5'-AAGCCATTGTGTCCGAACCA-3' | 5'-GAGCGGAAGAGCAGTTTGTCA-3' |
| | *Chat* | 5'-AGCCTTCCTAAGCCTCTACTG-3' | 5'-CTAAGCACACCAGAGATGAGG-3' |
| Schwann cell markers | *Ngf* | 5'-TTCCAGGCCCATGGTACAAT-3' | 5'-AAACTCCCCCATGTGGAAGAC-3' |
| | *Krox20* | 5'-TGCGCCTAGAAACCAGACCTT-3' | 5'-ATGCCCGCACTCACAATATTG-3' |
| | *Ncam-1* | 5'-TGGAACGCCGAGTACGAAGTA-3' | 5'-TGAACACGAAGTGAGCTGCCT-3' |
| | *Pmp22* | 5'-TGTACCACATCCGCCTTGG-3' | 5'-GAGCTGGCAGAAGAACAGGAAC-3' |

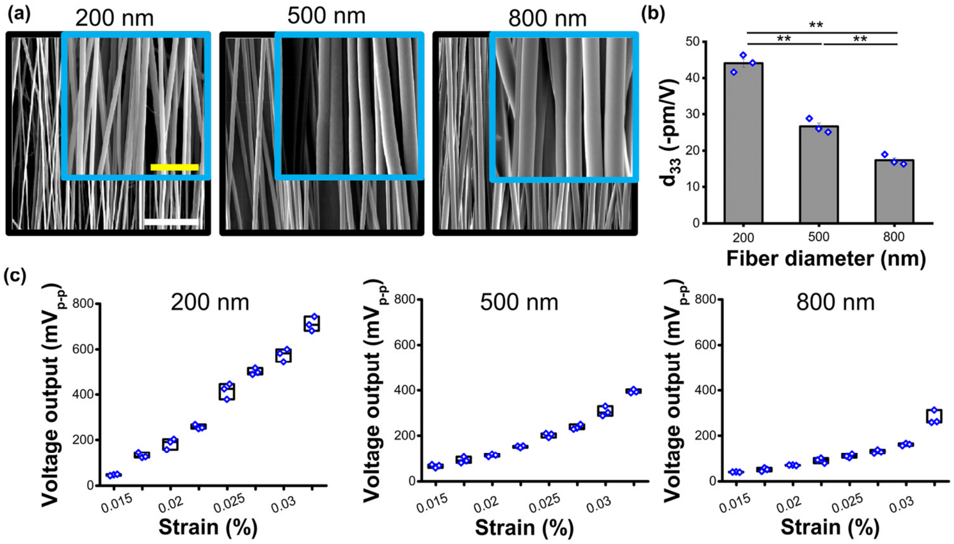


**Figure S1. Morphological and piezoelectrical characterization of electrospun P(VDF-TrFE) having various fiber diameters.** (a) Scanning electrospun microscopic (SEM) images of electrospun P(VDF-TrFE) scaffolds with an aligned fibrous structure having 200 nm, 500 nm, or 800 nm average fiber diameter (Yellow scale bar: 2 µm, While scale bar: 6 µm). (b) Piezo-force microscopy (PFM) measurements of individual electrospun P(VDF-TrFE) fibers having approximately 200, 500, or 800 fiber diameters. (c) The electric response of the electrospun P(VDF-TrFE) scaffolds with an average fiber diameter of 200, 500, or 800 nm (from left to right) under hydro-acoustic actuation that induces longitudinal strains from 0.015% to 0.0325%.

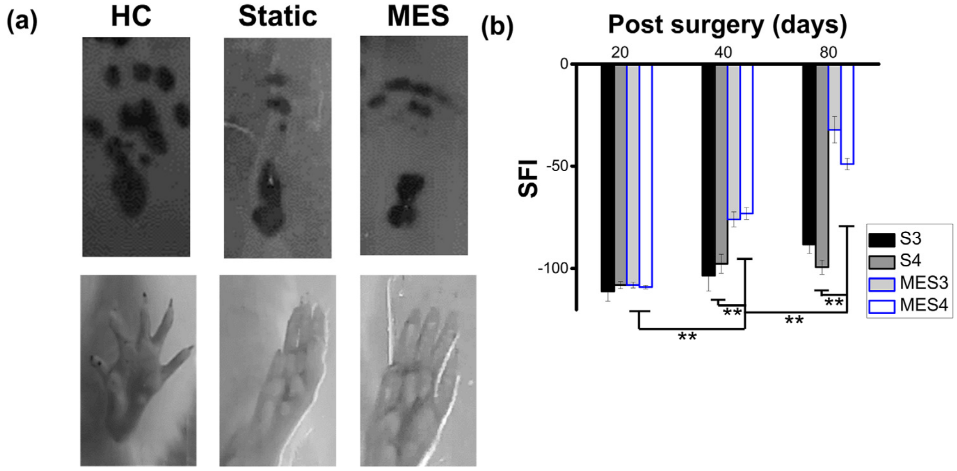


**Figure S2. Walking track analysis. Representative footprint images of walking track analysis and the corresponding sciatic functional index (SFI) quantification.** (a) Rat foot prints (top) collected for SFI measurements and photographs of rat paws (bottom) while the rats of healthy control (HC), static (Static) and MES conditions were walking through a transparent track. (b) Representative SFI data showing the functional motion recovery after the surgery at day 20, day 40, and day 80.

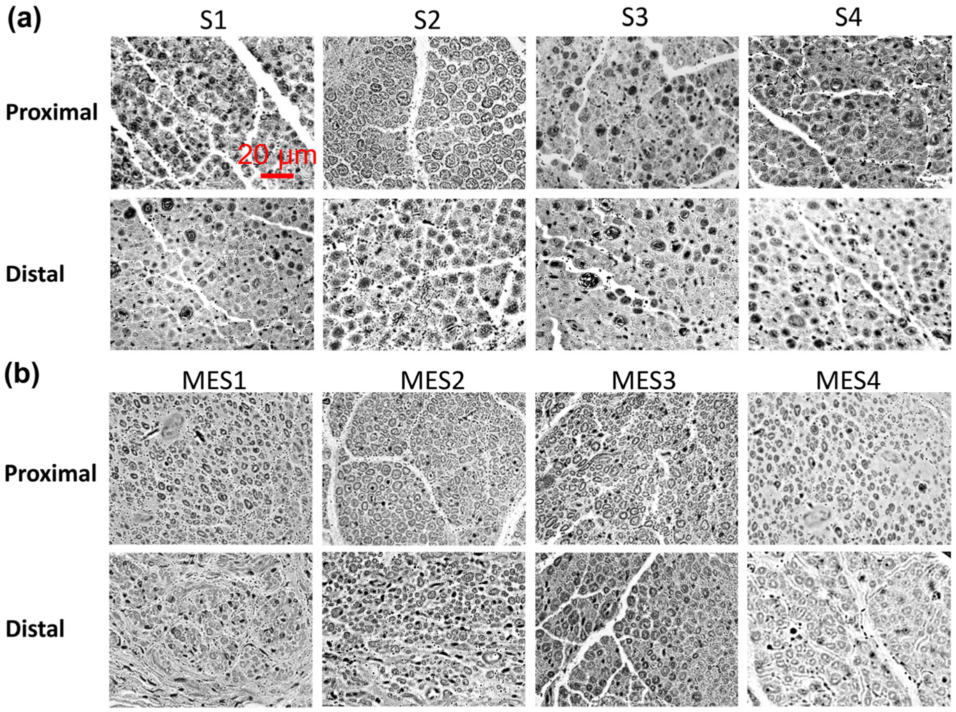


**Figure S3. Cross-sectional images of sciatic nerves (proximal and distal) in all experimental rats.** Cross-sectional images of proximal and distal ends of the 4 rats in (a) Static condition and (b) MES condition.

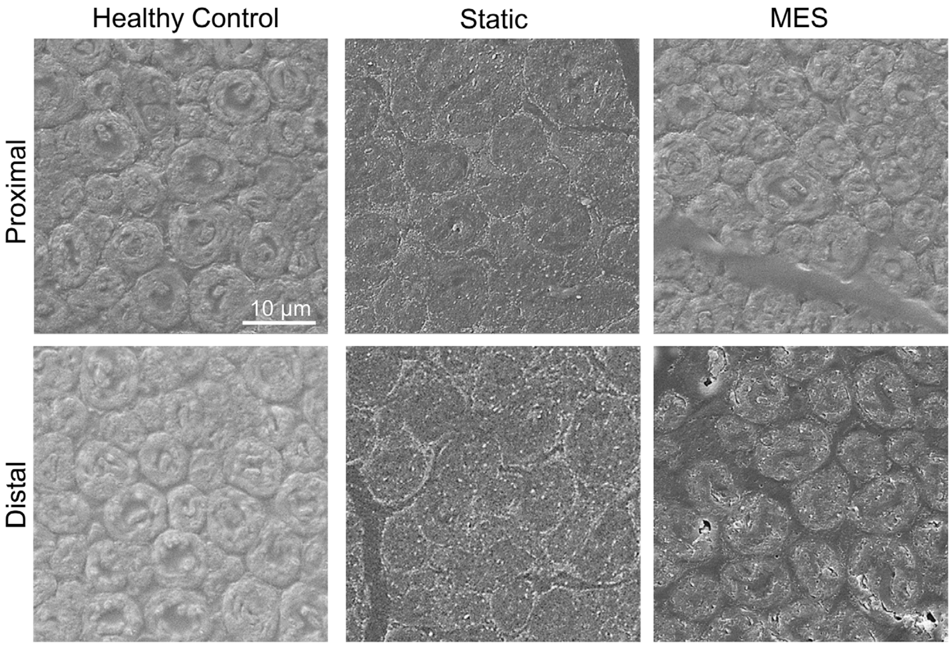


**Figure S4. Representative scanning electron microscopy (SEM) images of rat sciatic nerve.**

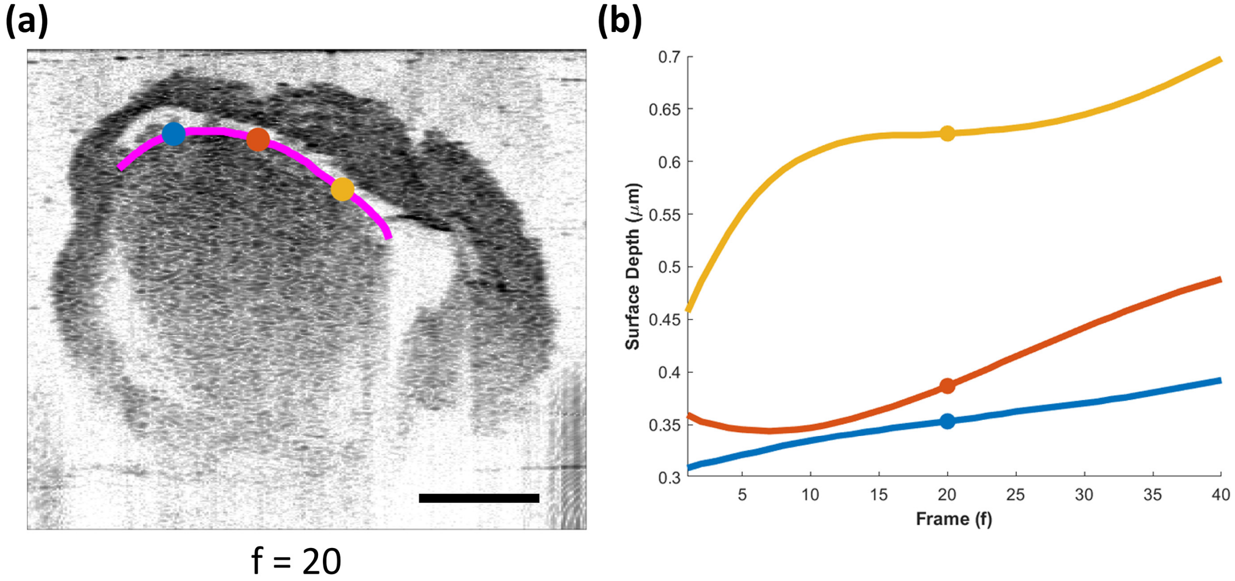


**Figure S5. Procedure for determining the nerve surface.** (a) The manually traced nerve surface (magenta line) is shown on a cross-sectional intensity image. In the intensity images, black and white color represent high and low back reflected light intensity, respectively. For ease of presentation, the depth at three distinct A-lines are plotted with blue, orange, and yellow dots. (b) The low-order polynomial fit of the depth values at different A-lines to obtain the nerve surface in every frame. Scale bar: 0.5 mm.

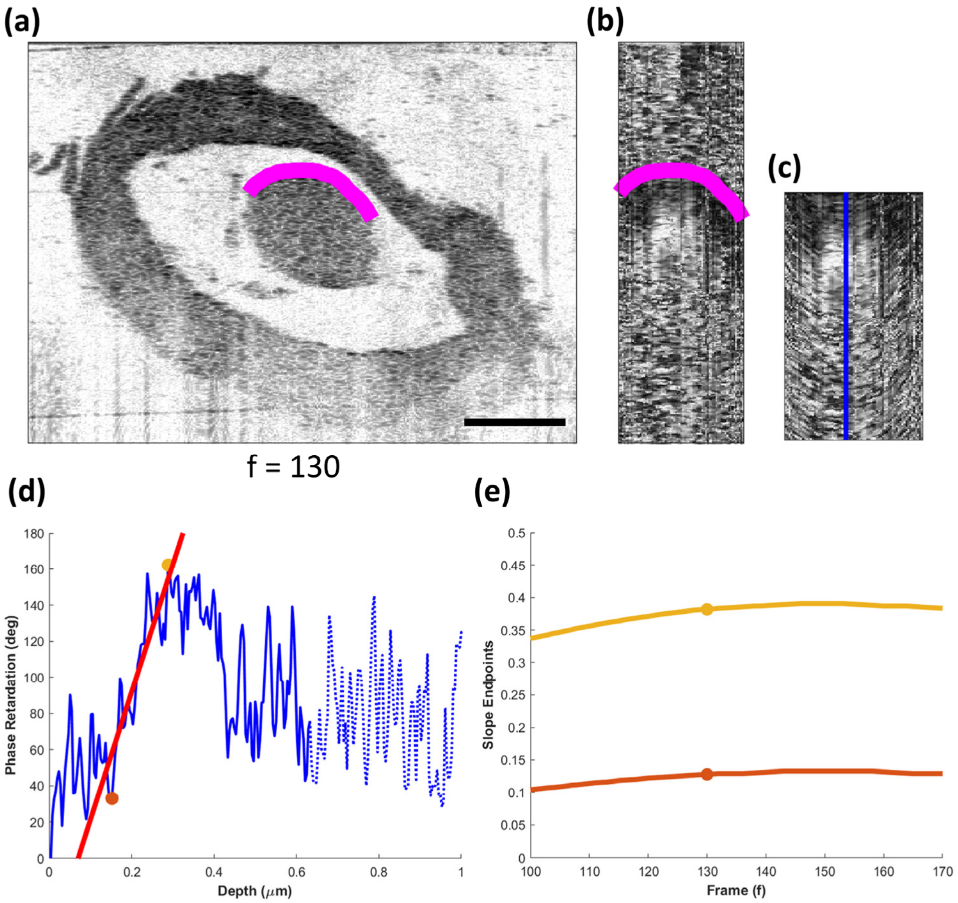


**Figure S6. Procedure for determining the endpoints for measuring the slope of the phase retardation curve.** (a, b) Cross-sectional intensity image and the corresponding cumulative phase retardation image of the nerve region. In the intensity image, black and white color represent high and low back reflected light intensity, respectively. In the phase retardation image, black and white represent 0 and 180 deg phase retardation, respectively. The magenta lines in (a) and (b) represent the surface of the nerve. (c) The height adjusted cumulative phase retardation image obtained by flattening the nerve surface in (b) so that the depth of the surface remains at 0. (d) The phase retardation vs depth curve taken from the A-line highlighted by the vertical blue line in (c). The orange and yellow dots in (d) denote the manually identified endpoints of the rising portion of the phase retardation curves. The broken lines in (d) highlights the regions where the degree of polarization uniformity is low (< 0.5). The red line in (d) is the linear least-square fit line from which the slope of the curve is obtained. (e) The endpoints across every ten frames in a volume section fit with a low-order polynomial to obtain the endpoints in every frame. Scale bar: 0.5 mm.

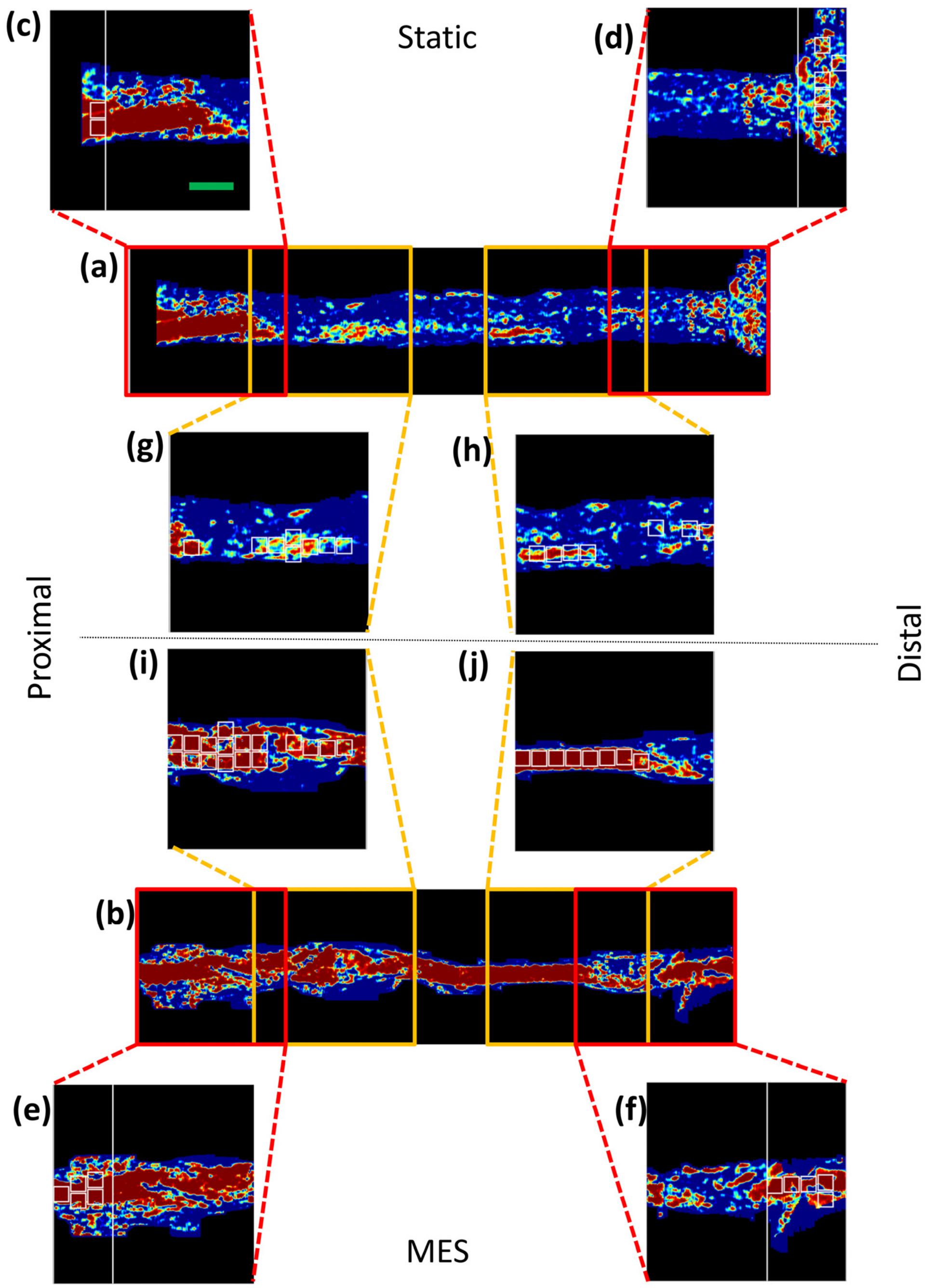


**Figure S7. Example images demonstrating the tiles for quantitative analysis.** (a) and (b) are the enface phase retardation slope images of the whole length of the samples S2 and MES2. In these images, blue to red color indicate low to high phase retardation in the range [0.07 0.2] deg $\mu m^{-1}$. The red rectangles in (a) and (b) highlight the volume sections used for quantitative comparison with the histological measurement. The yellow rectangles in (a) and (b) highlight the volume sections used for the quantitative comparison between the Static and MES groups. (c-j) shows the zoomed in images of the corresponding highlighted sections in (a) and (b). The vertical white lines in (c-f) denote the beginning/end of the conduit. The tiles in white rectangles in (c-j) highlight regions from where the average slopes were measured. The mean and standard error of these measurements across the tiles were calculated for both the proximal and distal ends for the different rats and were used for the quantitative comparison. Scale bar: 1 mm.
